## Extended Data Figures 1 to 15 for "The sugar-responsive enteroendocrine neuropeptide F regulates lipid metabolism through glucagon-like and insulin-like hormones in *Drosophila melanogaster*"

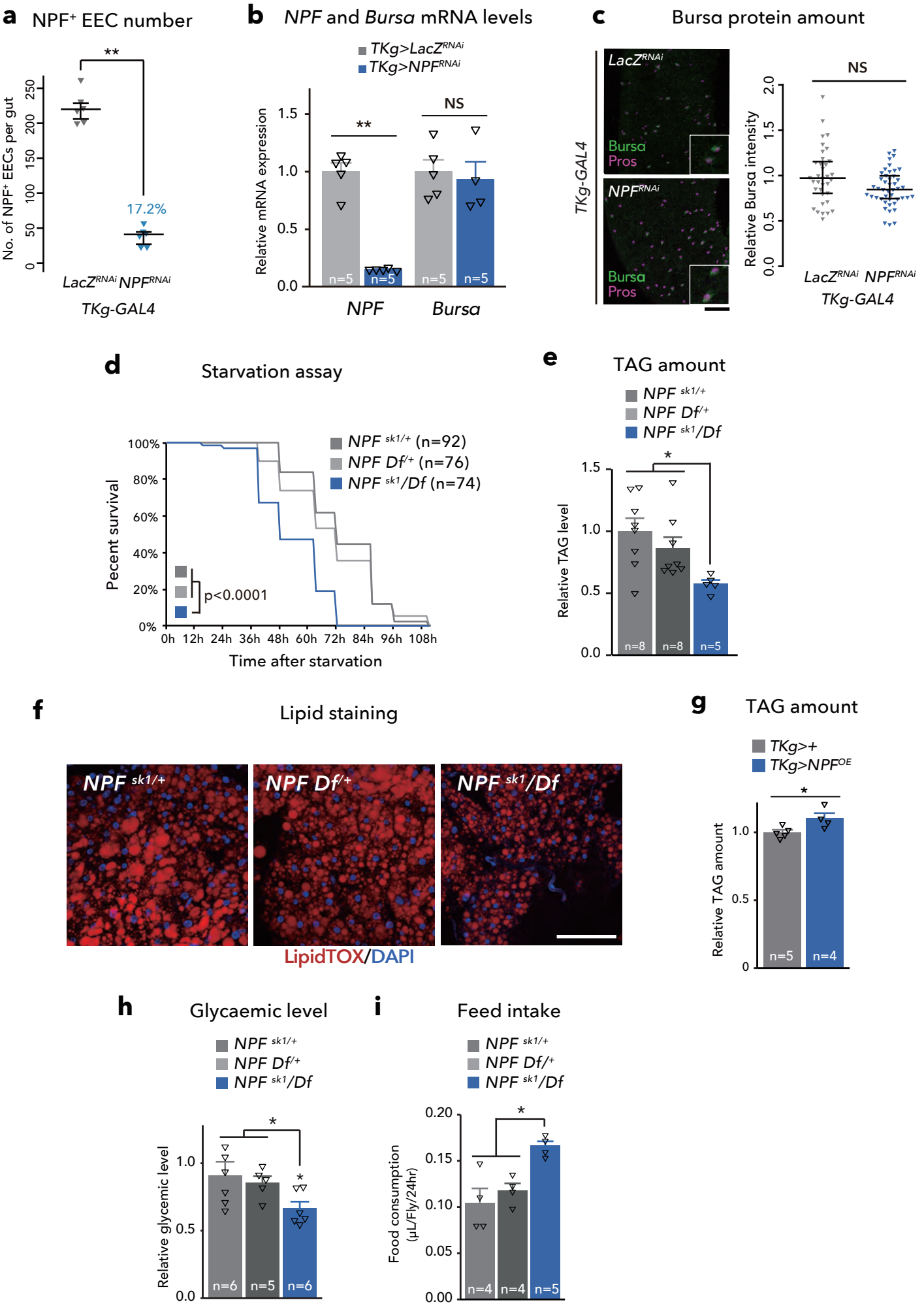

**Extended Data Figure 1. Midgut-specific *NPF* knockdown reduced the mRNA expression of *NPF*, but not *Bursa***

**a**, Number of *NPF*<sup>+</sup> EECs per midgut. n = 6 samples in each genotype. The three horizontal lines on each sample indicate lower, median, and upper quartiles. **b**, Relative change in the mRNA levels of neuropeptide genes in the gut. The number of samples assessed (n) is indicated in the graphs. **c**, Immunostaining for *Bursa* (green) and Prospero (magenta) and quantifications of *Bursa* fluorescent intensity in adult posterior midguts of control (*TKg>LacZ<sup>RNAi</sup>*) and *NPF* knockdown (*TKg>NPF<sup>RNAi</sup>*) animals. The number of EECs analysed in each genotype were more than 30. Each point represents *Bursa* fluorescent intensity in a single EEC. For each genotype, we used more than seven guts. Scale bar, 50 μm. **d**, Survival during starvation in flies of each genotype. The number of animals assessed (n) is indicated in the graphs. **e**, **g**, Relative TAG levels of each genotype. The number of samples assessed (n) is indicated in the graphs. **f**, LipidTOX (red) and DAPI (blue) staining of dissected fat body tissue of *NPF* mutant. Scale bar, 50 μm. **h**, Relative glycaemic level of *NPF* mutant animals. The number of samples assessed (n) is indicated in the graphs. **i**, Feeding amount measurement with CAFÉ assay. The number of samples assessed (n) is indicated in the graphs. For all bar graphs, mean and SEM with all data points are shown. Statistics: Wilcoxon rank sum test (a and c), two-tailed Student's t test (b, g), Log rank test (d), one-way ANOVA followed by Tukey's multiple comparisons test (e, h, and i). \*p < 0.05, \*\*p < 0.01; NS, non-significant (p > 0.05).

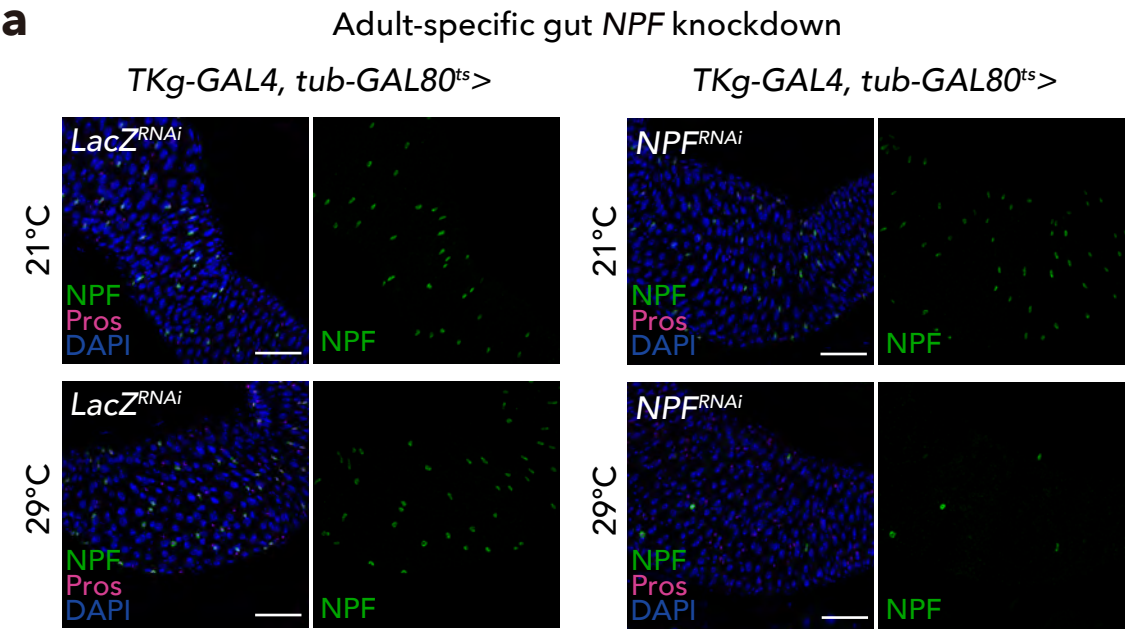

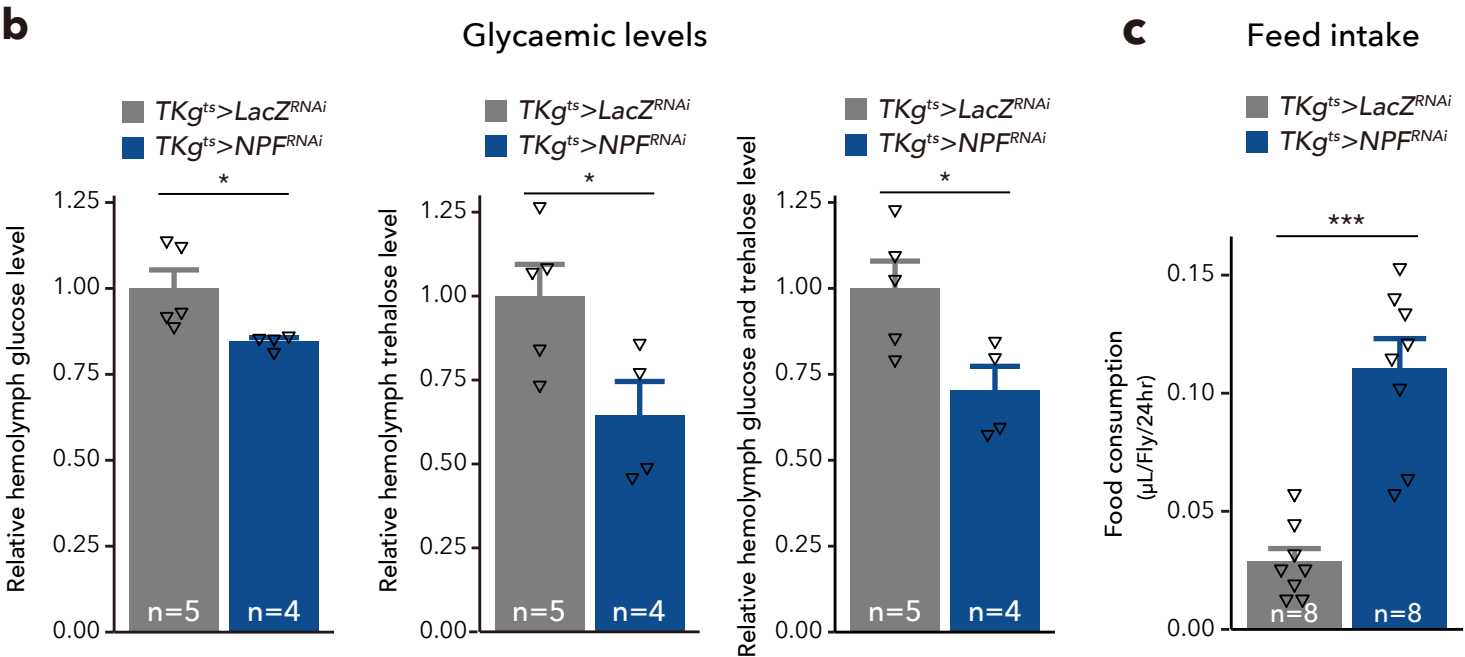

**Extended Data Figure 2. Adult-specific knockdown of *NPF* exhibited similar metabolic phenotypes to *TKg>NPF<sup>RNAi</sup>***

**a**, *TKg-GAL4*, *tub-GAL80<sup>ts</sup>*-mediated *NPF* knockdown (*TKg<sup>ts</sup>>NPF<sup>RNAi</sup>*) enabled the suppression of *NPF* expression in the gut only in the adult stage. Immunostaining for *NPF* (green), Prospero (marker for EEC), and DAPI (blue) in the adult middle midguts. **b**, Relative circulating levels of glucose (left), trehalose (mid), and the sum of glucose and trehalose (right) in *TKg<sup>ts</sup>>NPF<sup>RNAi</sup>* animals. The number of samples assessed (n) is indicated in each graph. **c**, Feeding amount measurement for each genotype with the CAFÉ assay. n = 8 samples, each point represents four adult female flies. For RNAi experiments, *LacZ* knockdown (*TKg<sup>ts</sup>>LacZ<sup>RNAi</sup>*) was used as negative control. For all bar graphs, mean and SEM with all data points are shown. Statistics: two-tailed Student's t test (b, and c), \*p < 0.05, \*\*p < 0.01, \*\*\*p < 0.001; NS, non-significant (p > 0.05).

Extended Data Figure3

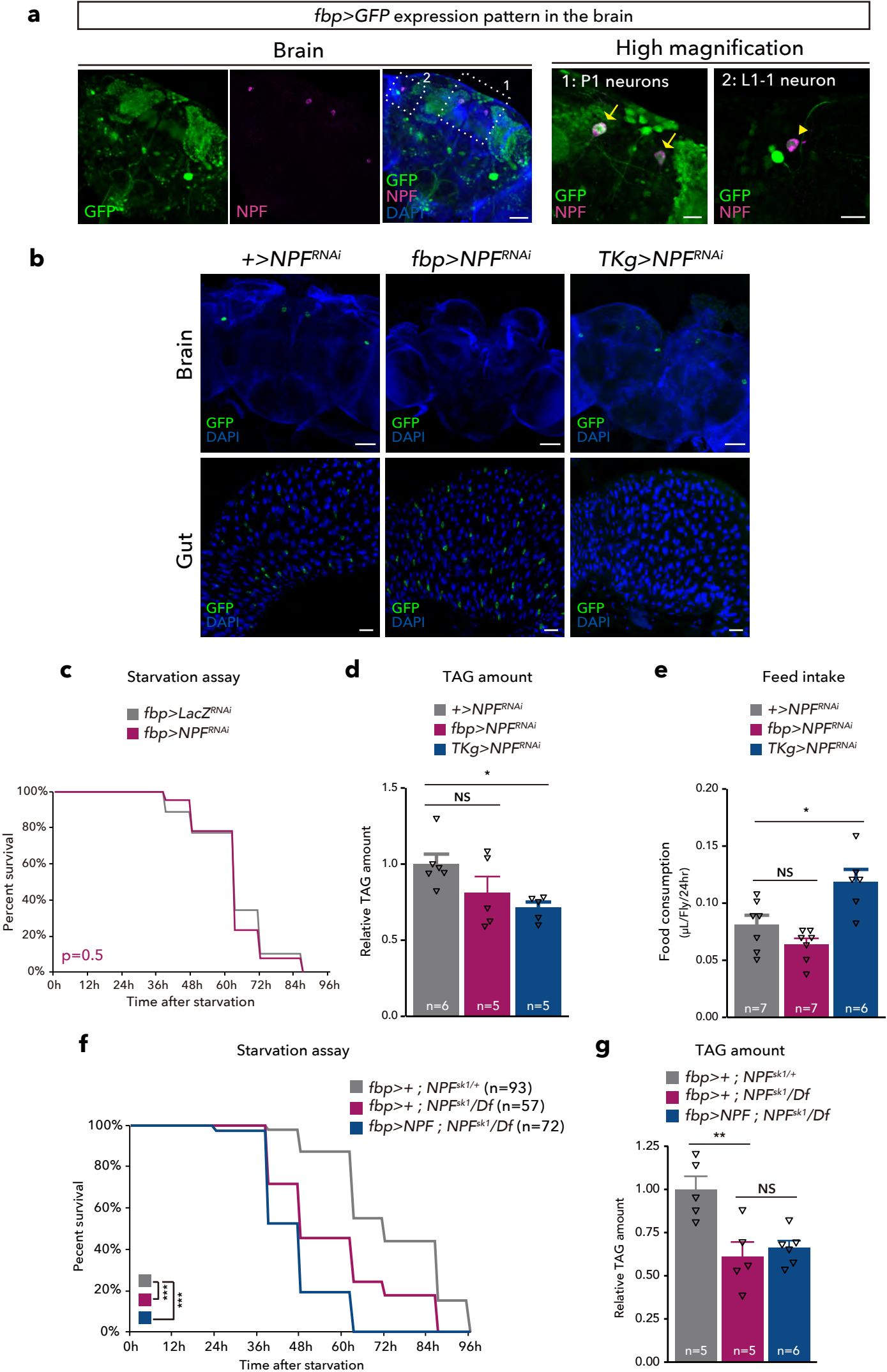

**Extended Data Figure 3. Knockdown of *NPF* in the brain does not reduce lipid storage.**

**a**, (left) Immunofluorescence of the brain in adult flies expressing *UAS-GFP* (green) reporter under *fbp-GAL4* (*fbp>GFP*). Cell bodies of NPF neurons are stained by anti-NPF antibody (magenta). Scale bar, 50 μm. (right) Magnified image of P1 NPF neurons (1), and L1-1 NPF neurons (2). Note that both NPF neurons were co-labelled with *fbp>GFP*. Scale bar, 20 μm. **b**, Immunostaining for NPF (green) and DAPI (blue) in adult brains (top), and gut (bottom) from control (*+>NPF<sup>RNAi</sup>*), brain-specific *NPF* knockdown animals (*fbp>NPF<sup>RNAi</sup>*), and gut-specific *NPF* knockdown animals (*TKg>NPF<sup>RNAi</sup>*). Scale bar, 50 μm. **c**, Survival during starvation in flies of control (*fbp>LacZ<sup>RNAi</sup>*) and *NPF* knockdown animals in the brain (*fbp>NPF<sup>RNAi</sup>*). The number of animals assessed (n) is indicated in each graph. **d**, Relative whole-body TAG levels of each genotype. The number of animals assessed (n) is indicated in each graph. **e**, Feeding amount measurement of each genotype with CAFÉ assay. The number of animals assessed (n) is indicated in each graph. Each point contains four adult female flies. **f**, Survival during starvation in flies of each genotype. The number of animals assessed (n) is indicated in each graph. **g**, Relative whole-body TAG levels of each genotype. The number of animals assessed (n) is indicated in each graph. For all bar graphs, mean and SEM with all data points are shown. Statistics: Log rank test with Holm's correction (c, f), one-way ANOVA followed by Tukey's multiple comparisons test (d, e, and g) \*$p < 0.05$, \*\*$p < 0.01$, \*\*\*$p < 0.001$; NS, non-significant ($p > 0.05$).

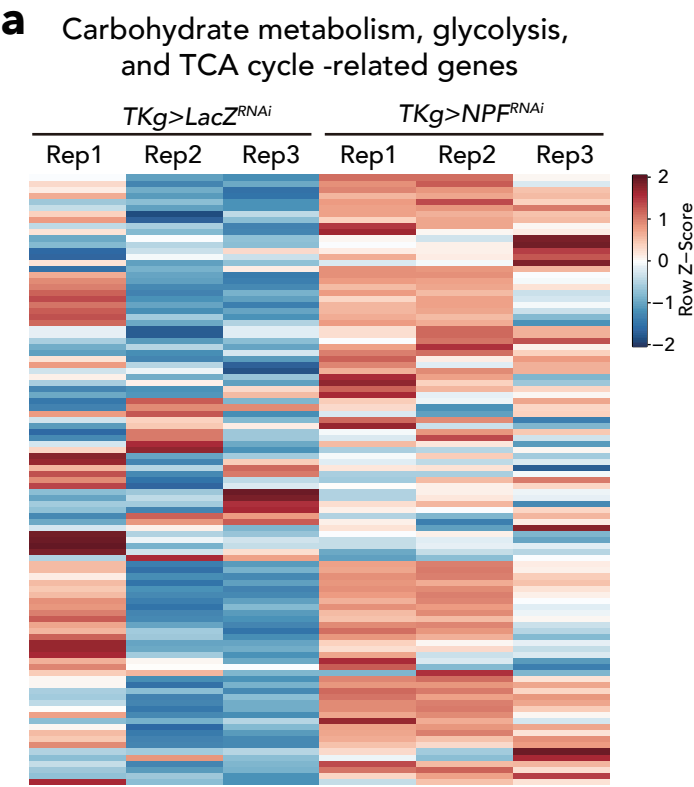

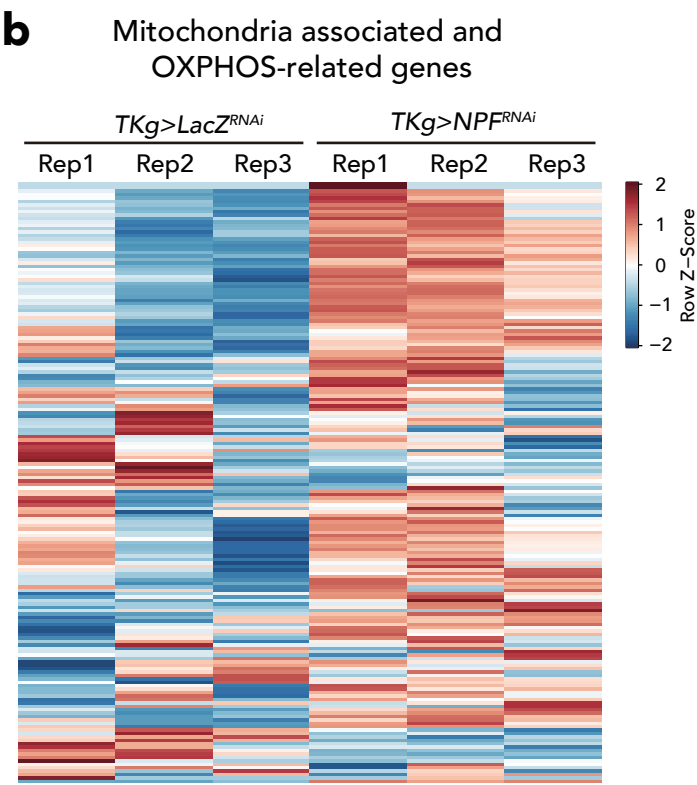

**Extended Data Figure 4. Knockdown of *NPF* in the gut changes carbohydrates and OXPHOS-related gene expression**

**a**, Expression heatmap of a curated set of metabolism genes showing trend of increase of carbohydrate metabolism, glycolysis, TCA cycle enzyme genes in *TKg>NPF<sup>RNAi</sup>* animals. Gene expression levels are represented by TMM-normalised FPKM. **b**, Expression heatmap of a curated set of mitochondria-associated genes showing trend of increase expression in *TKg>NPF<sup>RNAi</sup>* animals. Gene expression levels are represented by TMM-normalised FPKM.

**a** Cluster hetmap of whole body metabolomics data

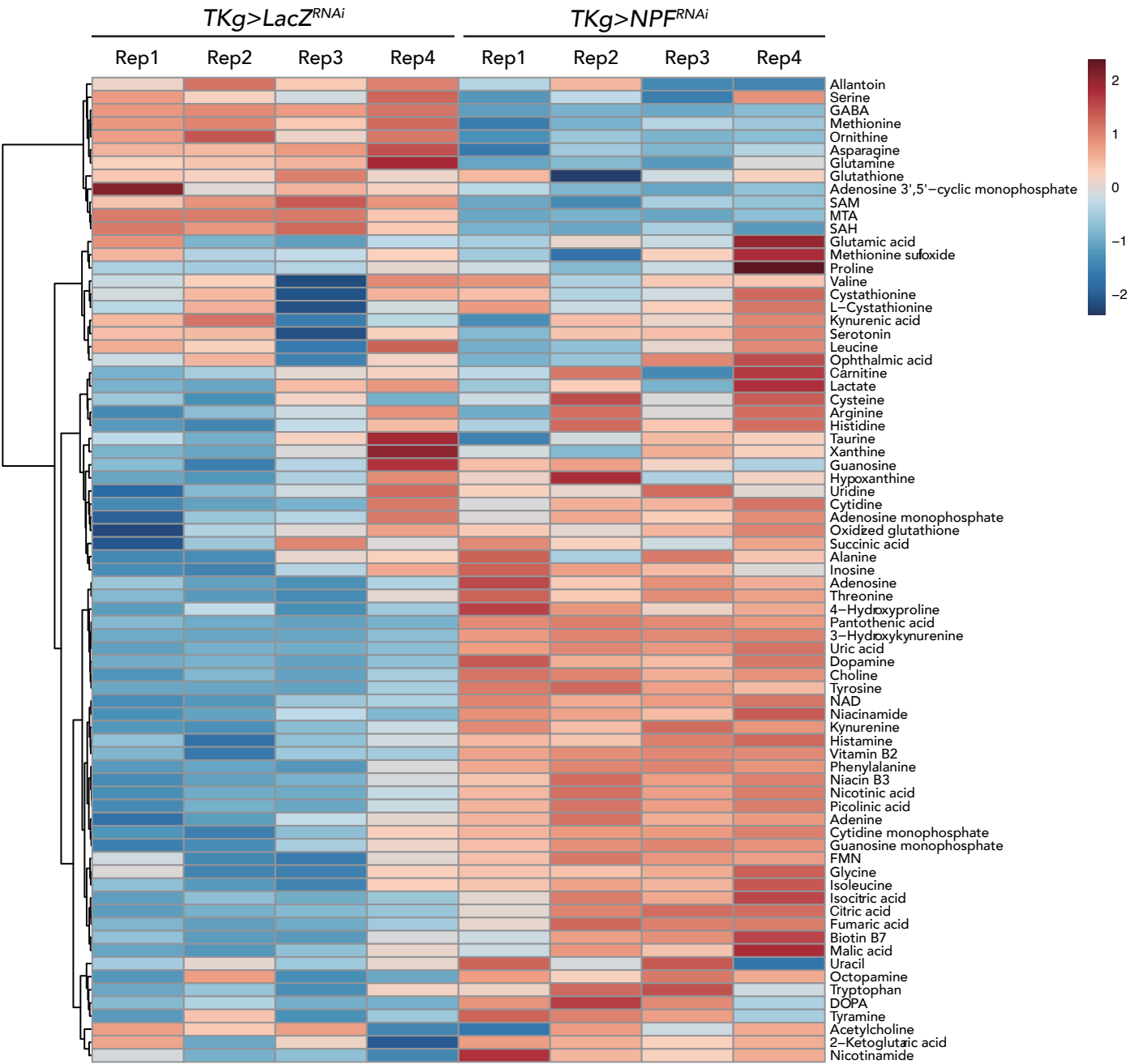

**Extended Data Figure 5. Knockdown of *NPF* in the gut changes metabolite profile of whole-body.**

**a,** Heatmap of clustering of changes of measured whole-body metabolite in *TKg>NPF<sup>RNAi</sup>* and *TKg>LacZ<sup>RNAi</sup>*. Note that NPF knockdown animals indicated dispersed cluster with control animals.

Extended Data Figure6

**a**

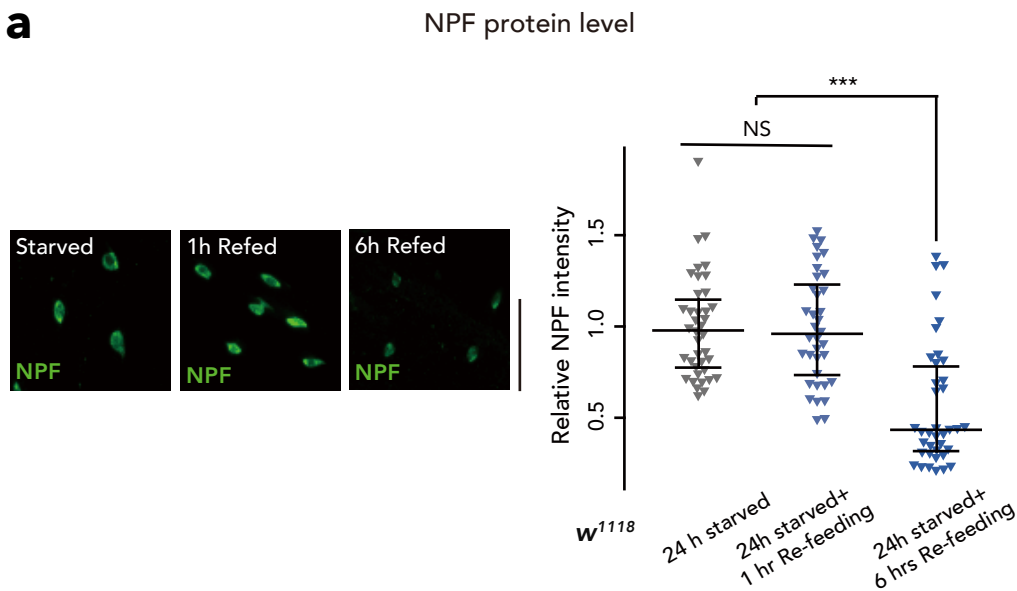

**b**

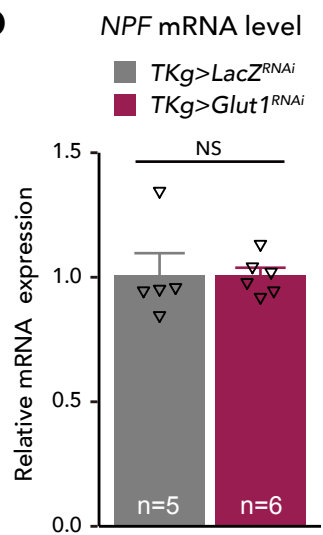

**c**

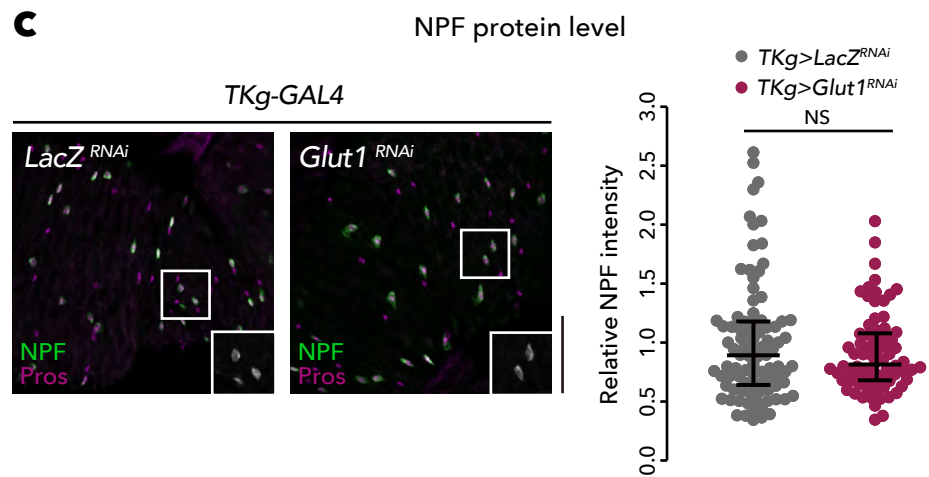

**d**

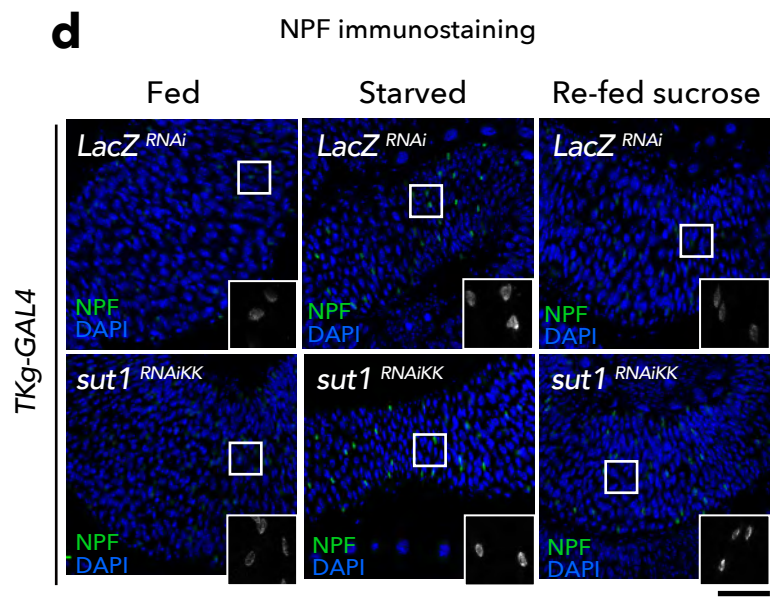

**e**

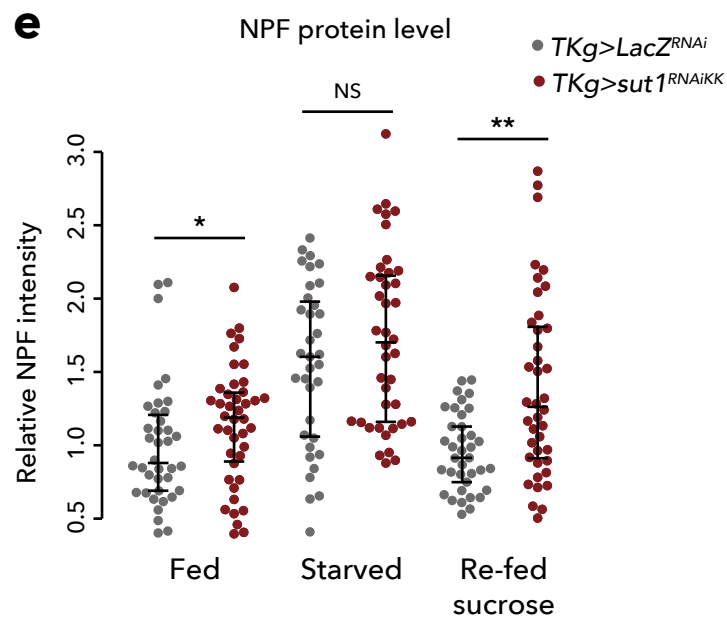

**Extended Data Figure 6. Midgut NPF accumulation is restored 6hre-feeding with sucrose**

**a**, (left) Immunostaining of NPF (green) in adult middle midguts from 6-day-old control ($w^{1118}$) animals starved for 24 h, and fed with sucrose for 1 or 6 h following a 24-h starvation. (right) Quantifications of NPF fluorescent intensity under the condition as in (a). $n > 30$, points of each condition were sampled with more than six guts and correspond to an individual EEC. Scale bar, 25 $\mu\text{m}$. **b**, RT-qPCR analysis of *NPF* mRNA level following *TKg-GAL4* mediated knockdown of *Glut1* ($TKg>Glut1^{RNAi}$). The number of samples assessed ($n$) is indicated in each graph. **c**, Immunostaining (left) and quantification of florescence (right) for NPF (green/white) and Prospero (magenta) in adult posterior midguts of control ($TKg>LacZ^{RNAi}$) and *Glut1* knockdown ($TKg>Glut1^{RNAi}$) animals. The number of EECs analysed in each genotype was greater than 35. Each point represents NPF fluorescent intensity in a single EEC. For each genotype, more than seven guts. Scale bar, 50 $\mu\text{m}$. **d**, Immunostaining for NPF (green/white) and DAPI (blue) in adult middle midguts from 6-day-old control $TKg>lacZ^{RNAi}$ and $TKg>sut1^{RNAi}$ animals fully fed (Fed), on 48 h starvation (Starved), and on re-feeding with sucrose following a 24 h starvation period (Re-fed sucrose). Scale bar, 50 $\mu\text{m}$. **e**, Quantifications of NPF fluorescent intensity of each genotype as in (d). The number of EECs analysed in each genotype was greater than 30. Each point represents NPF fluorescent intensity in a single EEC. For each genotype, more than eight guts were used. For all bar graphs, mean and SEM with all data points are shown. Statistics: Wilcoxon rank sum test with Holm' correction (a, c, and e), two-tailed Student's t test (b). \*$p < 0.05$, \*\*$p < 0.01$, \*\*\*$p < 0.001$; NS, non-significant ($p > 0.05$).

Extended Data Figure7

**a**

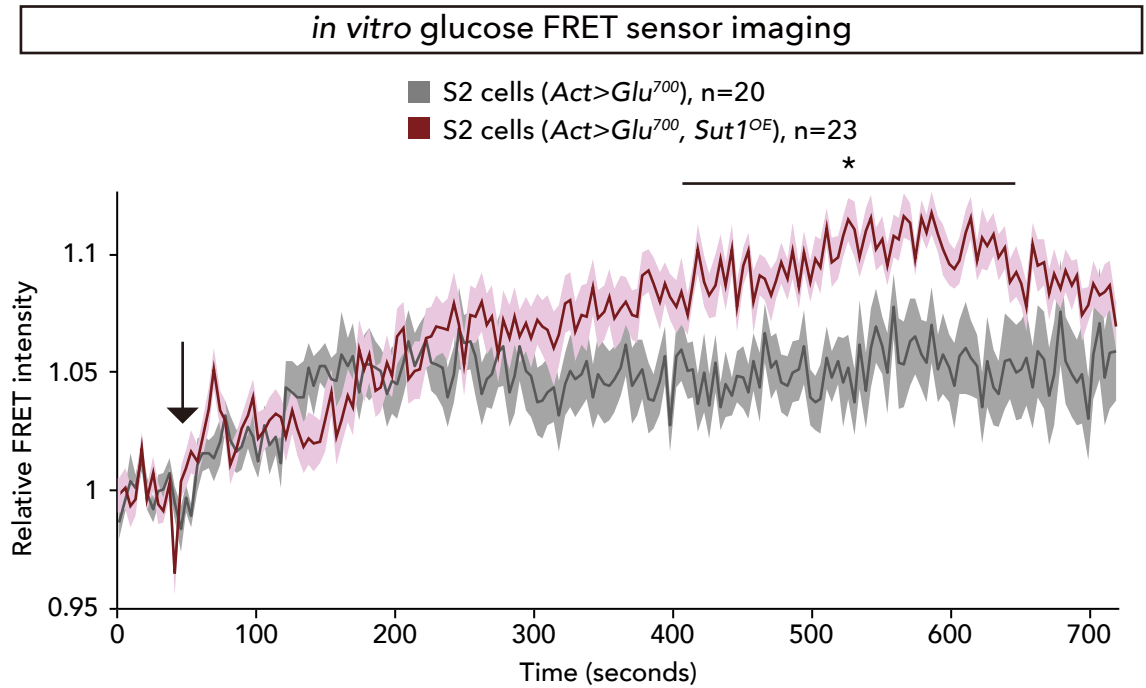

**b**

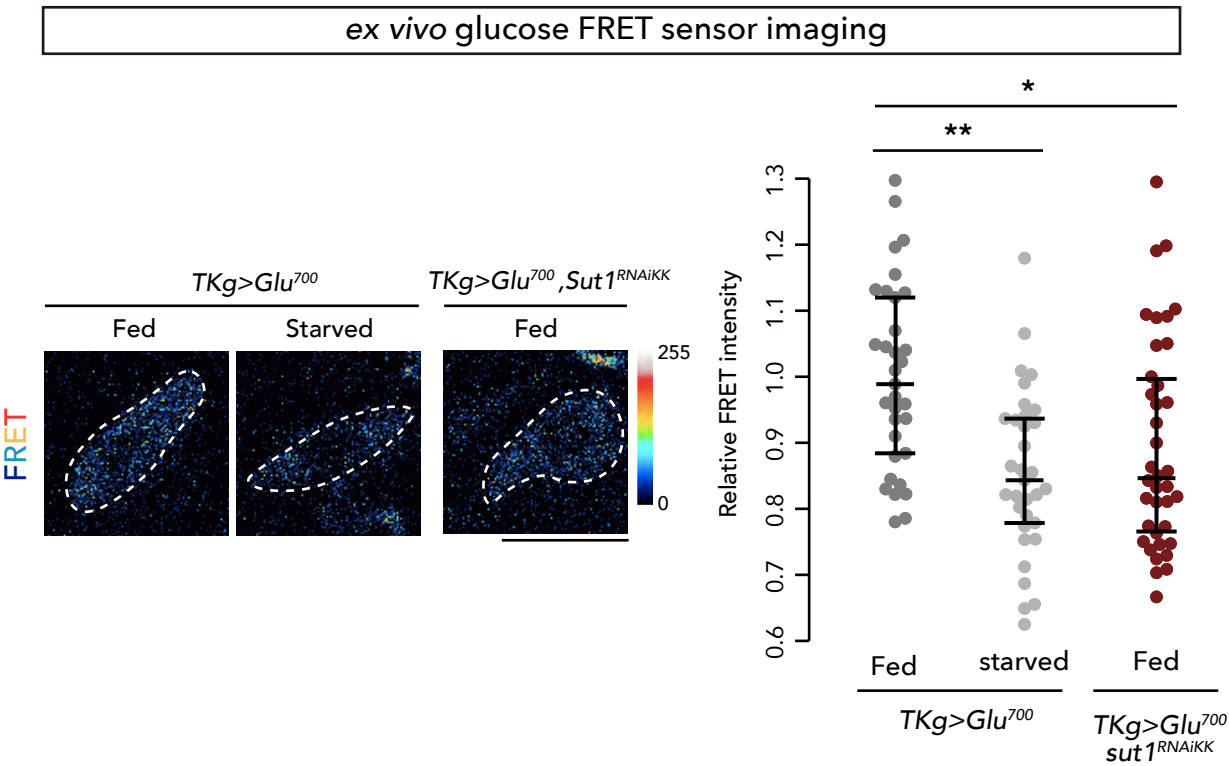

**Extended Data Figure 7. Sut1 in the EECs regulates intracellular glucose level**

**a**, Changes in the relative fluorescence intensity of Glu<sup>700</sup> glucose sensor FRET (YFP/CFP) signal of *Act>Glu<sup>700</sup>* S2 cells (grey), and *Act>Glu<sup>700</sup>, sut1<sup>OE</sup>* S2 cells (red) after 750 s exposure to glucose solution (final conc. 25 mM). Arrow indicates the time (40 s) of glucose administration. Note that *sut1* overexpression significantly increased the FRET signal in response to glucose administration between 400 s and 650 s. Statistical analysis was performed with average FRET levels from 400 s to 650 s of each genotypes. **b**, Glu<sup>700</sup> glucose sensor FRET signal in EECs of *ad libitum* feeding control (*TKg>Glu<sup>700</sup>*), 24 h starved control, and *ad libitum* feeding *sut1* knockdown (*TKg>Glu<sup>700</sup>, sut1<sup>RNAiKK</sup>*) animals. (Left) Representative FRET image in EECs (dashed lines). Scale bar: 10 μm. (Right) Quantification of FRET signals. Median FRET ratio for each genotype was set at 1 for *ad libitum* feeding control. Statistics: Wilcoxon rank sum test with Holm' correction (a, and b). \*p < 0.05, \*\*p < 0.01; NS, non-significant (p > 0.05).

Extended Data Figure8

*sut1* knockdown with *UAS-sut1<sup>RNAiTRiP</sup>* line

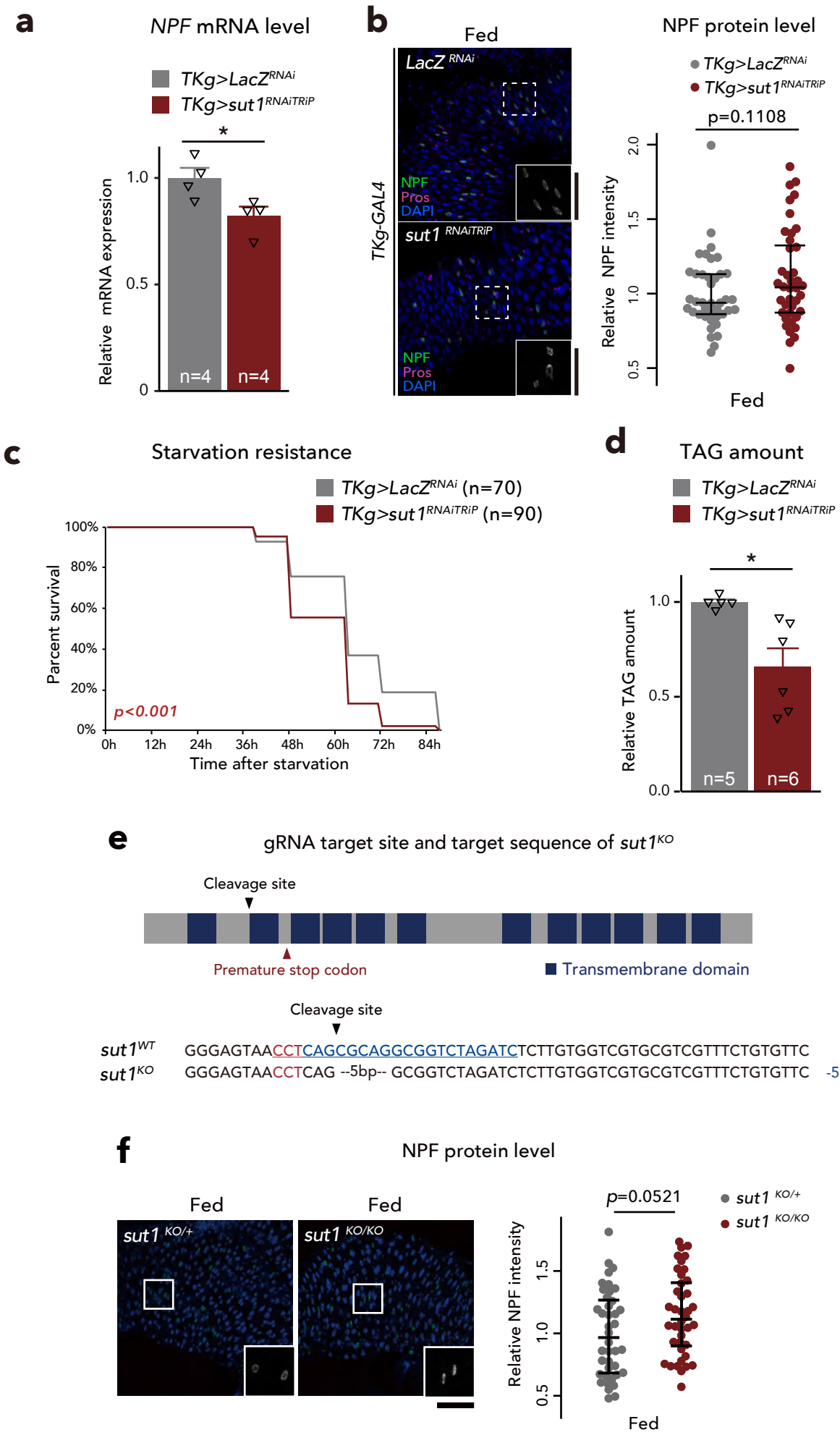

**Extended Data Figure 8. Sut1 in the EECs regulates NPF production**

**a**, RT-qPCR analysis of *NPF* mRNA level following *TKg-GAL4* mediated knockdown of *sut1* with another *UAS-sut1<sup>RNAi</sup>* line (*TKg>sut1<sup>RNAiTRiP</sup>*). The number of samples assessed (n) is indicated in the graph. **b**, (left) Immunostaining for NPF (green/white), Prospero (magenta), and DAPI (blue) in adult middle midguts from *TKg>sut1<sup>RNAiTRiP</sup>* animals. (right) Quantifications of NPF fluorescent intensity for each genotype. More than 30 EECs were analysed for each genotype. Each point represents NPF fluorescent intensity in a single EEC. For each genotype, more than eight guts were used. **c**, Survival during starvation in *TKg>sut1<sup>RNAiTRiP</sup>* flies. The number of animals assessed (n) is indicated in each graph. **d**, Relative whole-body TAG levels. The number of samples assessed (n) is indicated in each graph. **e**, A schematic representation of *sut1* coding sequence structure, gRNA target sequence (blue), and deletion of *sut1* knockout (*sut1<sup>KO</sup>*) mutant allele. Regions of the putative transmembrane domains of Sut1 are highlighted in dark blue. A premature stop codon generated in the *sut1<sup>KO</sup>* allele is indicated by a red arrowhead. DNA sequences of wild-type (WT) and *sut1<sup>KO</sup>* alleles are shown. The Cas9-gRNA target sequence is underlined. The PAM sequence is indicated in red. The 5 bp deletion results in a premature stop codon between the second and third transmembrane domains of Sut1. **f**, (left) Immunostaining for NPF (green/white) and DAPI (blue) in adult middle midguts from *sut1* mutant animals. Scale bar, 50 μm. (right) Quantification of NPF fluorescent intensity for each genotype as described in (e). The number of EECs analysed in each genotype was greater than 30. Each point represents NPF fluorescent intensity in a single EEC. For each genotype, more than eight guts were used. For RNAi experiments, *LacZ* knockdown (*TKg>LacZ<sup>RNAi</sup>*) was used as negative control. For all bar graphs, mean and SEM with all data points are shown. Statistics: two-tailed Student's t test (a, and d), Wilcoxon rank sum test (b, and f), Log rank test (c). \*p < 0.05; NS, non-significant (p > 0.05).

Extended Data Figure9

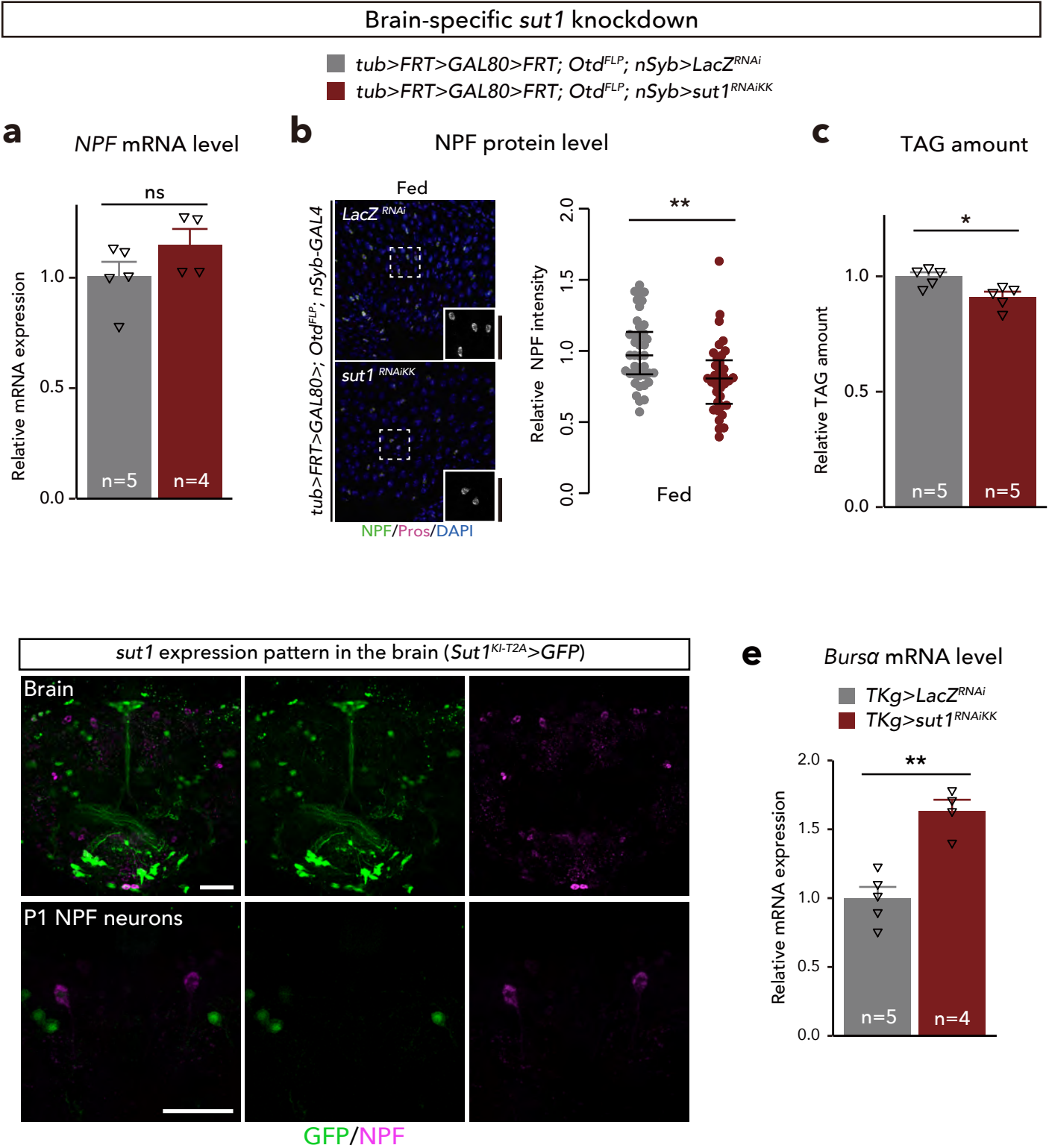

**Extended Data Figure 9. *sutI* knockdown in the brain does not attenuate gut NPF**

**a**, RT-qPCR analysis of *NPF* mRNA level in brain-specific *sutI* knockdown flies using *Otd-FLP* flies. The number of samples assessed (n) is indicated in the graph. **b**, (left) Immunostaining for NPF (green/white), Prospero (magenta), and DAPI (blue) in adult middle midguts from brain-specific *sutI* knockdown animals. (right) Quantification of NPF fluorescent intensity for each genotype. More than 30 EECs were analysed for each genotype. Each point represents NPF fluorescent intensity in a single EEC. For each genotype, more than eight guts were used. **c**, Relative whole-body TAG levels. The number of samples assessed (n) is indicated in each graph. **d**, Immunofluorescence of *sutI<sup>KI-T2A</sup>-GAL4*-driven *UAS-GFP* in the brain (top), and P1 NPF neurons (bottom) stained for GFP (green), and NPF (magenta). Note, no NPF<sup>+</sup> neurons exhibiting *sutI<sup>KI-T2A</sup>-GAL4*-driven GFP signals were detected. Scale bar, 50 μm. **e**, RT-qPCR analysis of *Bursa* mRNA level in *TKg>sutI<sup>RNAi</sup>* guts. The number of samples assessed (n) is indicated in the graph. For RNAi experiments, *LacZ* knockdown was used as negative control. For all bar graphs, mean and SEM with all data points are shown. Statistics: two-tailed Student's t test (a, c, and e), Wilcoxon rank sum test (b). \*p < 0.05, \*\*p < 0.01; NS, non-significant (p > 0.05).

Extended Data Figure10

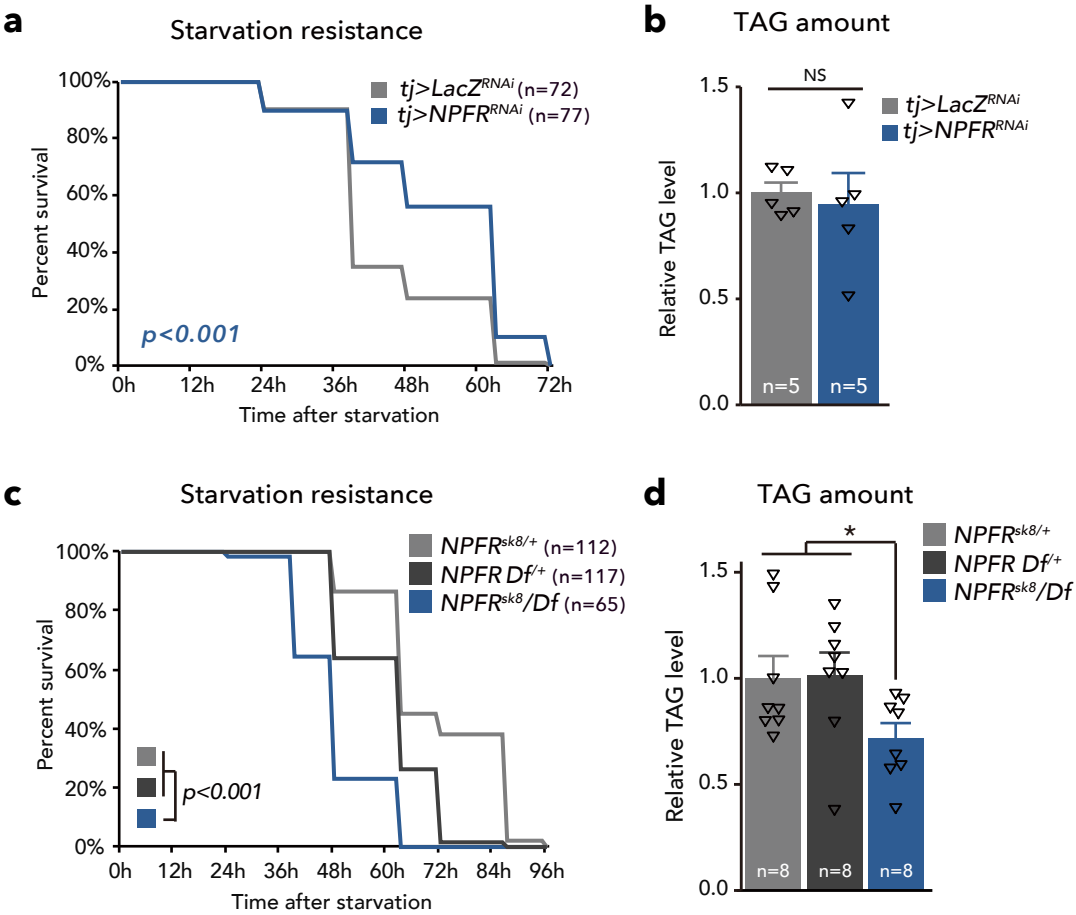

**Extended Data Figure 10. NPFR in the ovarian somatic cell does not affect lipid storage**

**a**, Survival during starvation in flies of control ($tj>LacZ^{RNAi}$) and *NPFR* knockdown animals in the ovarian somatic cells ($tj>NPFR^{RNAi}$). The number of animals assessed (n) is indicated in the graphs. **b**, Relative whole-body TAG levels of each genotype as in (a). The number of samples assessed (n) is indicated in the graphs. **c**, Survival during starvation in flies of control ($NPFR^{sk8/+}$ and $NPFR^{Df/+}$), and $NPFR^{sk8/Df}$. The number of animals assessed (n) is indicated in the graphs. **d**, Relative whole-body TAG levels of each genotype as in (d). The number of samples assessed (n) is indicated in the graphs. For all bar graphs, mean and SEM with all data points are shown. Statistics: Log rank test with Holm's correction (a, and c), two-tailed Student's t test (b), one-way ANOVA followed by Tukey's multiple comparisons test (d). \*p < 0.05; NS, non-significant (p > 0.05).

**a**

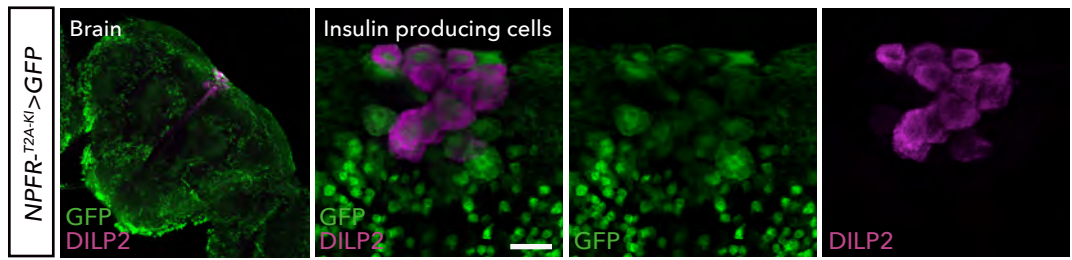

**b**

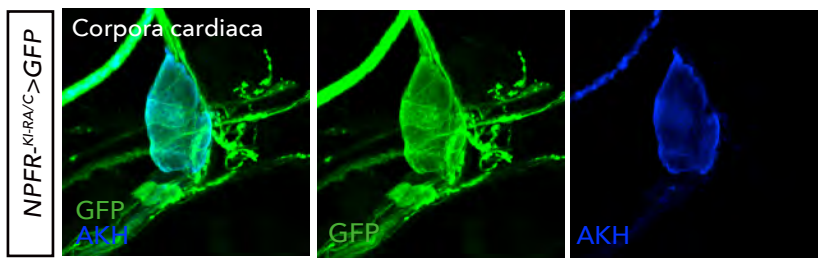

**c**

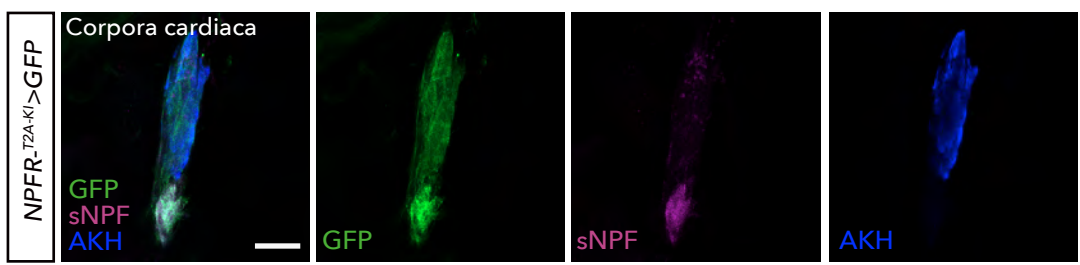

**d**

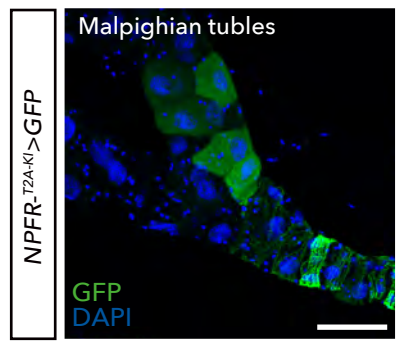

**e**

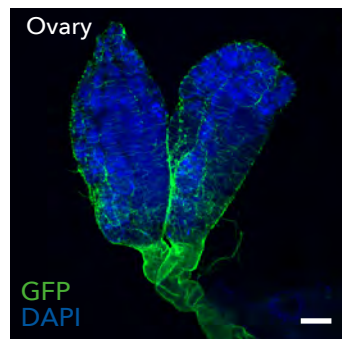

**f**

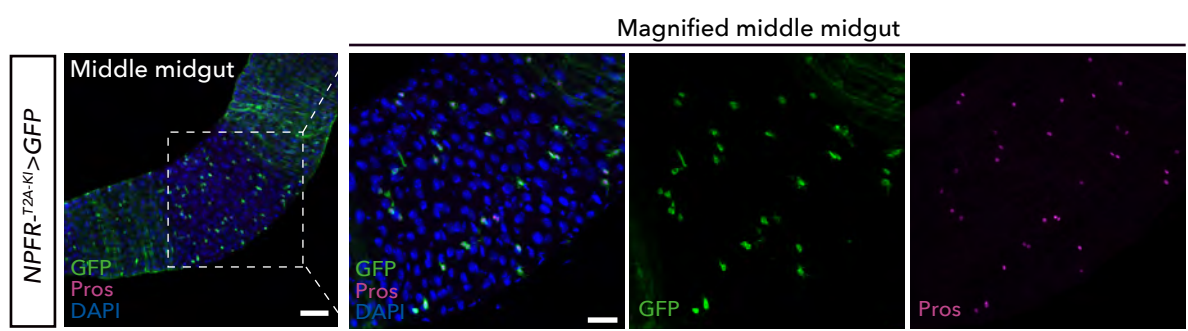

**Extended Data Figure 11. Expression pattern of *NPFR* knock-in *GAL4* lines**

**a**, Immunofluorescence of the brain in adult flies expressing *UAS- GFP* (green) reporter under *NPFR<sup>KI-T2A</sup>-GAL4*. Cell bodies of insulin-producing cells are stained by anti-DILP2 antibody (magenta). Scale bar, 20 μm. **b, c**, Immunofluorescence of the corpora cardiaca (CC) in adult flies expressing *UAS-GFP* (green) reporter under two *NPFR<sup>KI</sup>-GAL4* lines. Cell bodies of sNPF neurons and CC are stained by anti-sNPF antibody (magenta) and anti-AKH antibody (blue), respectively. Scale bar, 20 μm. **d**, Immunofluorescence of the Malpighian tubules in adult flies expressing *UAS- GFP* (green) reporter under *NPFR<sup>KI-T2A</sup>-GAL4*. Scale bar, 50 μm. **e**, Immunofluorescence of the ovary in adult flies expressing *UAS- GFP* (green) reporter under *NPFR<sup>KI-T2A</sup>-GAL4*. Scale bar, 100 μm. **e**, Immunofluorescence of the middle midgut in adult flies expressing *UAS- GFP* (green) reporter under *NPFR<sup>KI-T2A</sup>-GAL4*. Note that several GFP+ cells are labelled by the EEC marker Prospero (magenta). Scale bar, (left) 50 μm, (right) 25 μm.

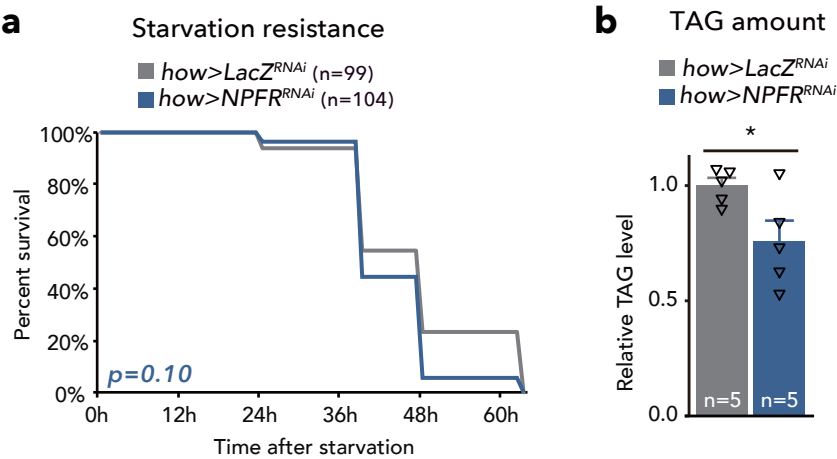

**Extended Data Figure 12. NPFR in the visceral muscle has small effect on starvation resistance and lipid amount**

**a**, Survival during starvation in flies of control (*how>LacZ<sup>RNAi</sup>*), and *NPFR* knockdown animals in the visceral muscle (*how>NPFR<sup>RNAi</sup>*). The number of animals assessed (n) is indicated in the graphs. **b**, Relative whole-body TAG levels of each genotype. The number of samples assessed (n) is indicated in the graphs. For bar graph, mean and SEM with all data points are shown. Statistics: Log rank test (g), two-tailed Student's t test (h). \*p < 0.05; NS, non-significant (p > 0.05).

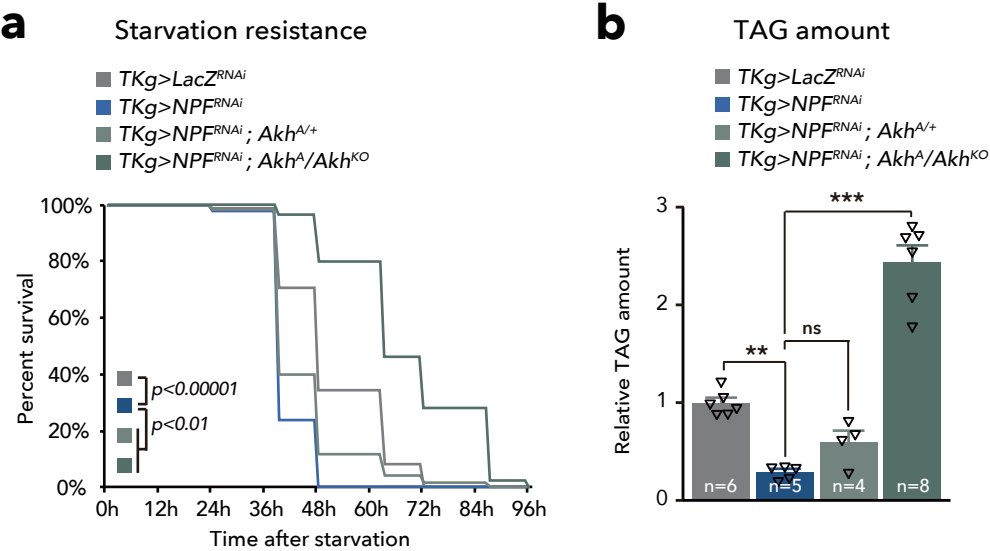

**Extended Data Figure 13. Loss of AKH restores starvation resistance and lipid reduction of *TKg>NPF<sup>RNAi</sup>***

**a**, Survival during starvation in flies of each genotype. The number of animals assessed (n) is indicated in the graphs. **b**, Relative whole-body TAG levels of each genotype. The number of animals assessed (n) is indicated in the graphs. For all bar graphs, mean and SEM with all data points are shown. Statistics: Log rank test with Holm's correction (a), one-way ANOVA followed by Tukey's multiple comparisons test (b). \*\*p < 0.01, \*\*\*p < 0.001; NS, non-significant (p > 0.05).

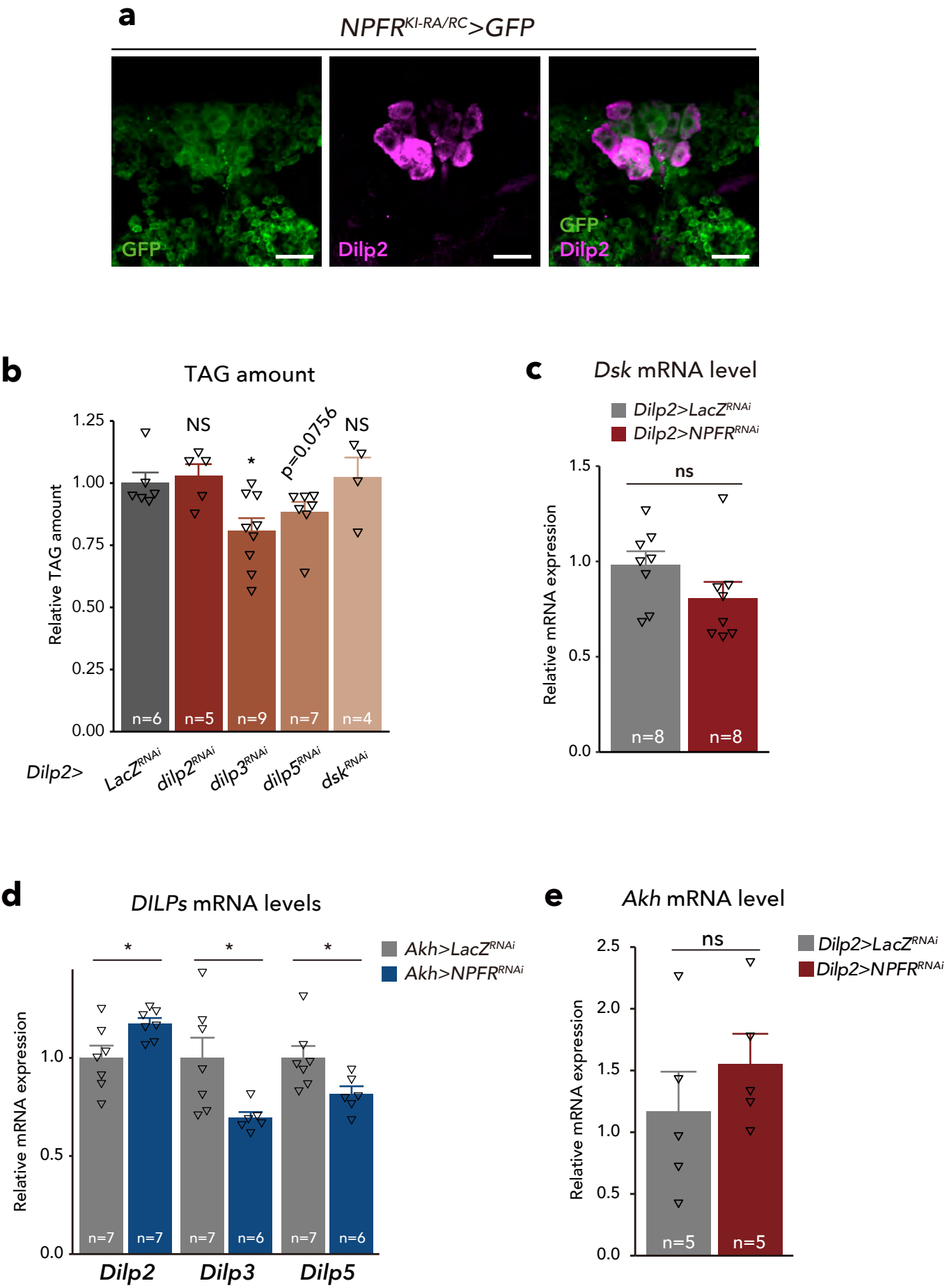

**Extended Data Figure 14. NPF neurons does not have direct connection with the IPCs**

**a**, Immunofluorescence of the IPCs in adult flies expressing *UAS-mCD8::GFP* (green) reporter under *NPFR<sup>KI-RA/RC</sup>-GAL4*. Cell bodies of IPCs are stained by anti-DILP2 (magenta). Scale bar, 20 μm. **b**, Relative whole-body TAG levels of each genotype. The number of samples assessed (n) is indicated in the graphs. **c**, RT-qPCR analysis of *Dsk* mRNA level in *Dilp2>NPFR<sup>RNAi</sup>* flies. The number of samples assessed (n) is indicated in the graph. **d**, RT-qPCR analysis of *dilps* mRNA level in CC-specific *NPFR* knockdown flies (*Akh>NPFR<sup>RNAi</sup>*). The number of samples assessed (n) is indicated in the graph. **e**, RT-qPCR analysis of *Akh* mRNA level in IPC-specific *NPFR* knockdown flies (*Dilp2>NPFR<sup>RNAi</sup>*). The number of samples assessed (n) is indicated in the graph.

For RNAi experiments, *LacZ* knockdown was used as negative control. For all bar graphs, mean and SEM with all data points are shown. Statistics: two-tailed Student's t test (b-e). \*p < 0.05; NS, non-significant (p > 0.05).

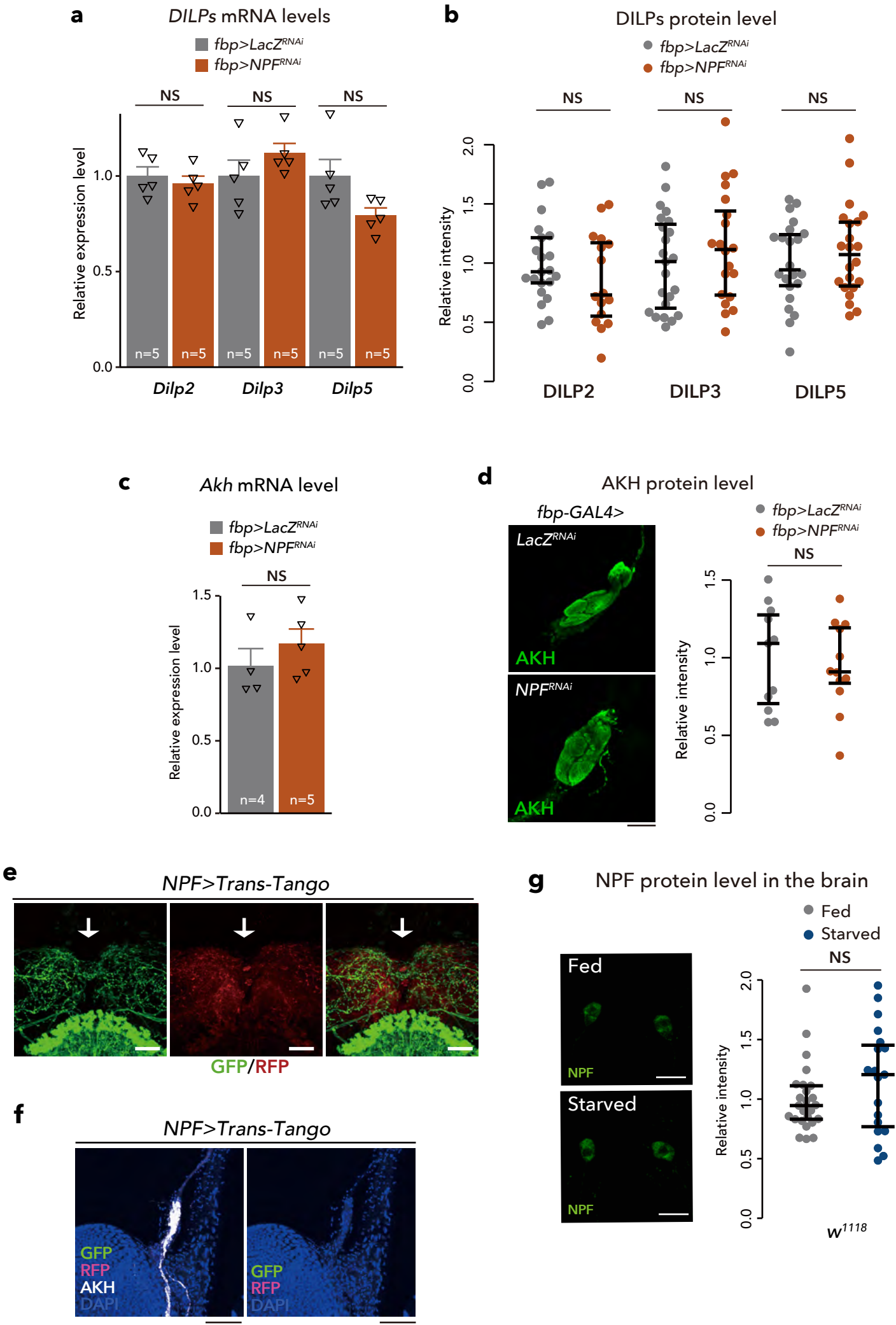

**Extended Data Figure 15. Brain NPF does not affect the levels of Dilps and AKH**

**a**, RT-qPCR analysis of *dilps* mRNA level in brain-specific *NPF* knockdown flies (*fbp>NPF<sup>RNAi</sup>*). The number of samples assessed (n) is indicated in the graph. **b**, Quantification of DILP2, 3, and 5 in the brain of adult *fbp>NPF<sup>RNAi</sup>* animals. Scale bar, 20 μm. n > 20. **c**, RT-qPCR analysis of *Akh* mRNA levels in the whole bodies of *fbp>NPF<sup>RNAi</sup>* adult animals. The number of samples assessed (n) is indicated in the graph. **d**, Quantification of AKH (green) in the CC of *fbp>NPF<sup>RNAi</sup>* adult flies. Scale bar, 10 μm. n > 10. **e**, Immunofluorescence of the PI region (arrow) in adult flies expressing *Trans-Tango* driven by *NPF-GAL4* (*NPF>Trans-Tango*). Scale bar, 20 μm. Note, postsynaptic signal (RFP; red) was not observed in the PI region. **f**, Immunofluorescence of the CC in *NPF>Trans-Tango* adult flies. Cell bodies of CC are stained with anti-AKH antibody (white). Scale bar, 10 μm. Note, neither presynaptic signals (GFP; green) nor postsynaptic signals (RFP; magenta) were observed near the CC. **g**, (left) Immunostaining for NPF (green) of P1 NPF neurons in the adult brains of *ad libitum* feeding, and 24h starved *w<sup>1118</sup>* flies. Scale bar, 20 μm. (right) Quantification of NPF fluorescent intensity. n > 15. For RNAi experiments, *LacZ* knockdown (*fbp>lacZ<sup>RNAi</sup>*) was used as negative control. For all bar graphs, mean and SEM with all data points are shown. Statistics: two-tailed Student's t test (a, and c), Wilcoxon rank sum test (b, d, and g), NS, non-significant (p > 0.05).
